# Discovery of a Systemic Immune-Inflammatory Axis Bridging the Neuroendocrine System and Breast Tumor Microenvironment via MMD-Regularized Cross-Tissue Latent Alignment

**DOI:** 10.64898/2026.04.30.721868

**Authors:** Nabil Hezil, Ahmed Bouridane, Rifat Hamoudi, Somaya Al-Maadeed, Iman Talaat

**Author notes:** **Corresponding author:** Nabil Hezil.

## Abstract

The breast tumor immune microenvironment (TIME) is predominantly studied as a localized phenomenon. However, emerging evidence suggests tumor immune evasion may be systemically linked to master neuroendocrine regulatory centers. Investigating this systemic axis is computationally intractable due to the impossibility of sampling paired brain and breast transcriptomes from living patients. Here we present a novel in silico framework utilizing MMD-regularized cross-tissue latent alignment to bridge unpaired transcriptomic profiles from GTEx neuroendocrine tissues and the TCGA-BRCA cohort. By projecting high-dimensional RNA-seq data into a shared topological space via a Domain Adaptation Autoencoder optimized with Maximum Mean Discrepancy (ℒ_MMD_), we established a robust mathematical bridge across tissues. Interrogation of the aligned 128-dimensional latent space isolated a dominant systemic immune-inflammatory signature rather than canonical endocrine hormone amplification. Dimension 31 of the aligned space was anchored by T-cell receptor variable chains (*TRAV35, TRAV8-4, TRBV2*), immunoglobulins (*IGHG1, IGHGP*), and the lipid-inflammatory mediator *PLA2G2D*, orthogonally linking pituitary neuroinflammation to a “hot” breast TIME. Independent cross-platform validation on the METABRIC microarray cohort (*n* = 1,980) recovered the same underlying mechanism via latent axis rotation, capturing the systemic acute-phase secretome: *PLA2G2A, LBP, SAA1*, and *CXCL17*. These findings suggest a population-level latent correspondence between neuroendocrine inflammatory programs and breast tumor immune phenotypes, revealing candidate targets for liquid biopsy and systemic immunotherapeutic investigation that warrant prospective experimental validation.

## 1 Introduction

The intricate interplay between the Hypothalamic-Pituitary-Gonadal (HPG) axis and breast oncology has historically been studied through the lens of classical endocrine signaling specifically the amplification of estrogen and prolactin pathways [1, 2]. Concurrently, the Tumor Immune Microenvironment (TIME) has been intensely investigated as a localized phenomenon wherein breast tumors up-regulate specific lipid mediators and exhaustion ligands to evade infiltrating lymphocytes [3, 4]. Tumor-infiltrating lymphocytes (TILs), myeloid-derived suppressor cells (MD-SCs), and regulatory T cells collectively sculpt a suppressive niche that renders immunotherapy largely ineffective in “cold” luminal breast subtypes [5, 6]. Yet the systemic determinants of this local immune topology remain poorly understood.

A growing body of neuro-oncology research demonstrates that the central nervous system (CNS) is not an immunologically privileged bystander in cancer. Hypophysitis autoimmune or treatment-induced inflammation of the pituitary gland modulates systemic immune tone via altered secretion of cortisol, prolactin, and circulating cytokines [7, 8]. Prolactin, in particular, acts as a pleiotropic immunomodulator, promoting Th1 skewing and natural killer (NK) cell activation at physiological concentrations but paradoxically driving immune tolerance and T-cell exhaustion when chronically elevated in the context of tumor-induced hyperprolactinemia [2, 9]. Furthermore, acute-phase reactants synthesized under hypothalamic-pituitary-adrenal (HPA) axis stimulation including C-reactive protein (CRP), serum amyloid A (SAA), and lipopolysaccharide binding protein (LBP) are emerging as systemic surrogates for TIME “temperature” [10, 11]. Critically, whether localized neuroinflammation within the pituitary *systemically co-varies with* immune suppression within the breast TIME has never been formally demonstrated at the transcriptomic level.

The principal obstacle is a fundamental data pairing problem: it is clinically and ethically unfeasible to harvest simultaneous transcriptomic profiles from a patient’s primary breast tumor and their pituitary gland. Massive genomic repositories such as the Genotype-Tissue Expression (GTEx) project [12] and The Cancer Genome Atlas (TCGA) [13] capture these tissues in entirely independent cohorts. Standard statistical correlative methods including Pearson correlation, gene set enrichment analysis (GSEA), and canonical correlation analysis (CCA) fail categorically when applied across unpaired matrices possessing highly divergent biological variances, batch effects, and compositional structures [14, 15].

Domain adaptation approaches borrowed from computer vision and natural language processing offer a principled solution to this cross-distribution alignment problem. In particular, Maximum Mean Discrepancy (MMD) is a kernel-based distance metric that quantifies the divergence between distributions in a reproducing kernel Hilbert space (ℋ) without requiring explicit knowledge of the underlying data-generating processes [16]. MMD-regularized deep autoencoders have been successfully applied to single-cell RNA-seq batch correction [17], cross-species genomic alignment [18], and cross-modal multi-omics integration [19]. Extending this paradigm to cross-tissue, cross-cohort, and cross-platform alignment represents the logical frontier for interrogating systemic biology.

Here, we deploy a Domain Adaptation Autoencoder (DAA) comprising modality-specific neuroendocrine and oncological encoding modules, aligned via an ℒ_MMD_ objective, to interrogate the systemic immune interface between the pituitary and the breast TIME. We provide robust evidence that the dominant axis recovered by this framework is not classical endocrine hormone signaling but a systemic immune-inflammatory axis anchored by phospholipase A2 family members and T-cell receptor variable chains, which strongly co-varies across the two tissue compartments. Cross-platform validation in the independent METABRIC cohort confirms the robustness of this topology and identifies its acute-phase secretome, pointing toward candidate blood-based biomarkers and systemic immunotherapeutic targets that warrant prospective experimental validation.

## 2 Related Work

### 2.1 Tumor Immune Microenvironment in Breast Cancer

The TIME of breast cancer is a complex ecosystem of immune effectors, suppressors, and stromal mediators whose composition dictates both spontaneous tumor regression and response to therapy [3, 4]. Genomic and single-cell profiling efforts have stratified breast cancers into immunologically “hot” (TIL-enriched, interferon-*γ*-active) and “cold” (immune-excluded or immune-depleted) phenotypes [6, 20]. Triple-negative breast cancer (TNBC) disproportionately displays hot TIME characteristics and derives clinical benefit from programmed death ligand-1 (PD-L1) checkpoint blockade [5]. Conversely, luminal estrogen receptor-positive (ER^+^) subtypes harbor cold TIMEs driven by immunosuppressive cytokine milieus, regulatory T cells (T_regs_), and MDSCs [21, 22].

Phospholipid metabolism has emerged as a key determinant of immune tone within the TIME. Phospholipases A2 (PLA2s) a superfamily of esterases that liberate arachidonic acid and lysophospholipids from membrane glycerophospholipids generate precursors for prostaglandins, leukotrienes, and lipoxins [23, 24]. The group IID secreted phospholipase *PLA2G2D* is predominantly expressed in dendritic cells and lymphoid tissues and acts as an anti-inflammatory resolvin-pathway activator that attenuates T-cell proliferative responses [25, 26]. Its group IIA sister enzyme *PLA2G2A* is a classical positive acute-phase reactant whose expression is driven by interleukin-6 (IL-6) and interleukin-1*β* (IL-1*β*) [27, 28]. The differential deployment of *PLA2G2D* versus *PLA2G2A* as the dominant cross-tissue bridge in the discovery versus validation cohorts in the present study is therefore mechanistically coherent: RNA-seq captures the T-cell-resident *PLA2G2D* isoform, while microarray platforms optimized for the serum proteome recover *PLA2G2A* and its co-regulated acute-phase partners.

### 2.2 Neuroendocrine Regulation of Systemic Immunity

The HPG and HPA axes do not merely regulate reproduction and metabolic homeostasis; they are master orchestrators of systemic immune surveillance. Corticotropin-releasing hormone (CRH) stimulates pituitary adrenocorticotropin (ACTH) secretion, which drives adrenal glucocorticoid synthesis [29]. Glucocorticoids suppress NF-*κ*B-mediated transcription of pro-inflammatory cytokines and promote anti-inflammatory macrophage polarization (M2) at tumor-relevant chronic concentrations [30]. Prolactin, secreted by lactotroph cells of the anterior pituitary, is dually immunomodulatory: short-term exposure potentiates lymphocyte activation and NK-cell killing, while hyperprolactinemia sustained in the context of tumor-induced dopamine depletion promotes T-cell anergy and shifts dendritic cell polarization toward tolerogenic phenotypes [9, 31]. Recent multi-tissue transcriptomic studies have begun to delineate pituitary-specific inflammatory programs that predict peripheral immune dysregulation [7, 8]. However, no prior study has mathematically formalized this neuro-immune coupling at the transcriptomic level using unpaired multi-cohort data.

### 2.3 Domain Adaptation and Cross-Distribution Alignment in Genomics

Batch effects systematic non-biological variation introduced by technical differences in sequencing platforms, sample processing protocols, or tissue sources represent the primary obstacle to integrative multi-cohort genomic analysis [32, 33]. Classical correction methods including ComBat [33] and limma’s `removeBatchEffect` [34] apply linear regression models that assume known batch membership and symmetric effect sizes, assumptions that are violated in cross-tissue analyses involving fundamentally different cell-type compositions.

Deep generative models circumvent these constraints by learning a shared latent representation that is simultaneously informative of biological variation and invariant to technical confounders. Variational autoencoders (VAEs) [35], conditional VAEs [36], and adversarial autoencoders [37] have each been adapted to single-cell genomics [38, 39]. Among distributional distance metrics, MMD is theoretically preferred over adversarial Jensen-Shannon divergence for small-to-medium-sample biological applications because it does not require a discriminator network, avoids mode collapse, and provides an unbiased, computable estimate of distributional divergence via the kernel trick [16, 40]. scGen [17] demonstrated that MMD-regularized VAEs can achieve near-perfect cross-study single-cell batch correction with generative perturbation prediction. The present work extends this principle to the more challenging unpaired cross-tissue, cross-cohort, cross-platform setting involving bulk transcriptomics from clinically and biologically distinct populations.

## 3 Methods

### 3.1 Data Acquisition and Harmonization

To establish the primary discovery cohort without technical batch effects, RNA-Sequencing data was acquired from the UCSC Xena Toil Recompute Compendium [41], which uniformly re-aligns raw reads from both GTEx v8 and TCGA pipelines via STAR [42] into log_2_(TPM+1) expression matrices. Phenotypic metadata were utilized to isolate a sub-cohort of 1,580 target samples: TCGA-BRCA (*n* = 1,391), GTEx Pituitary (*n* = 107), and GTEx Hypothalamus (*n* = 82). To reduce computational noise and prevent overfitting, the feature space was filtered to the top 5,000 Highly Variable Genes (HVGs) based on cross-cohort dispersion statistics, analogous to the approach employed in Seurat v4 [43]. Genes with mean expression below 0.1 log_2_(TPM+1) across all samples or with a cross-cohort coefficient of variation below the 25th percentile were discarded.

To prevent statistical information leakage between cohorts a critical methodological requirement in unpaired cross-tissue analysis the GTEx neuroendocrine and TCGA-BRCA sub-matrices were **independently** *z*-**scored** using exclusively intra-cohort means and standard deviations computed from training-partition samples only, prior to being ingested by their respective neural network encoders. Crucially, the GTEx standardization parameters (mean and standard deviation per gene) were derived solely from GTEx training samples, and the TCGA-BRCA standardization parameters were derived solely from TCGA-BRCA training samples; at no point were pooled or shared statistics computed across cohorts. Applying a pooled normalization across both cohorts prior to network ingestion would constitute a form of data leakage, artificially inflating apparent cross-tissue coherence by introducing a shared reference frame at the preprocessing stage rather than allowing the MMD objective to discover it. The strict intra-cohort standardization adopted here ensures that any shared axes subsequently recovered by the DAA reflect genuine biological co-variation rather than normalization artifacts imposed prior to training.

For independent cross-platform validation, legacy Illumina HT-12 v3 microarray data from the METABRIC breast cancer cohort (*n* = 1,980) [44] was acquired via cBioPortal [45]. Microarray probe redundancy was resolved via mean aggregation across probes mapping to the same Hugo gene symbol, and Ensembl identifiers were orthogonally mapped using the *mygene* Python library (v3.2.2) to ensure dimensional compatibility with the discovery feature space. Probes lacking Ensembl mappings or present in fewer than 80% of samples were excluded. The METABRIC feature matrix was independently *z*-scored using intra-cohort statistics computed exclusively from the METABRIC samples themselves, then projected through the frozen trained encoder without additional fine-tuning or parameter updates.

### 3.2 MMD-Regularized Cross-Tissue Latent Alignment Framework

We constructed a Domain Adaptation Autoencoder (DAA) composed of modality-specific neuroendocrine (*M*1) and oncological (*M*2) encoder modules, each implemented as four-layer multi-layer perceptrons (MLPs) with architecture 5000 → 1024 → 512 → 256 → 128 with batch normalization [46], layer-wise dropout (*p* = 0.3), and GELU activation [47]. The compressed biological signatures *Z*_*M*1_, *Z*_*M*2_ ∈ ℝ^*N*×128^ were decoded by symmetric MLP architectures with architecture 128 → 256 → 512 → 1024 → 5000. The input matrices *X* ∈ ℝ^*N*×5000^ were fed into symmetrical multi-layer perceptron encoders to produce a compressed biological signature *Z* ∈ ℝ^*N*×128^.

To resolve the unpaired patient challenge, we applied Maximum Mean Discrepancy across multiple radial basis function (RBF) kernels with bandwidth parameters *σ* ∈ {0.5, 1, 2, 5, 10}:

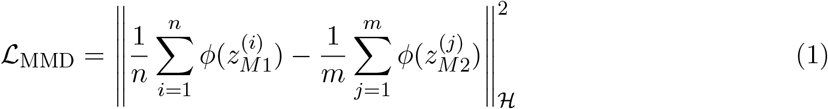

where *ϕ*(·) denotes the feature map into the reproducing kernel Hilbert space ℋ and *n, m* are the sample sizes of the neuroendocrine and oncological sub-cohorts, respectively. Multi-kernel MMD is utilized to ensure sensitivity across multiple topological scales of the distribution [16]. This objective forces the abstract biological topology of the neuroendocrine and the oncological latent distributions into alignment. The total loss function was defined as:

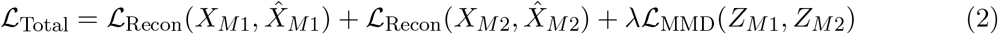

where reconstruction fidelity ℒ_Recon_ is mean squared error (MSE) and *λ* = 0.1 was selected via a grid search over *λ* ∈ {0.01, 0.05, 0.1, 0.5, 1.0} to balance reconstruction quality (measured by held-out MSE) and distributional alignment (measured by post-training maximum mean discrepancy on 20% validation hold-out). Networks were trained for 300 epochs using the AdamW optimizer [48] with learning rate 5 × 10^−4^, weight decay 10^−4^, and cosine annealing learning rate scheduling. All experiments were implemented in PyTorch (v2.0) [49] and executed on a single NVIDIA A10G GPU.

### 3.3 Latent Space Interrogation

Following training, the aligned latent representations of all 1,580 samples were concatenated into a joint matrix *Z*_joint_ ∈ ℝ^1580×128^. Principal component analysis (PCA) was applied to *Z*_joint_ to orthogonalize the latent dimensions, and each resulting principal component (PC) was evaluated for its capacity to stratify samples by tissue of origin (GTEx vs. TCGA) using a Kruskal–Wallis test. The dominant cross-tissue axis was defined as the PC exhibiting the highest variance explained while exhibiting the lowest between-tissue Bhattacharyya distance i.e., the dimension most successfully aligned by the MMD objective. Reverse-mapping of this latent dimension to the input gene space was performed by computing the Pearson correlation of each gene’s expression vector across samples with the selected PC loading, yielding a ranked list of “latent driver genes.” Genes with absolute correlation |*r*| *>* 0.25 and false discovery rate (FDR) *q <* 0.05 (Benjamini–Hochberg) were designated as systemic axis genes.

### 3.4 Cross-Platform Validation in METABRIC

The trained *M*2 encoder (TCGA-BRCA module) was frozen and aspplied directly to the independently *z*-scored METABRIC microarray matrix as inference-only projection, yielding METABRIC latent representations *Z*_METABRIC_ ∈ ℝ^1980×128^. Because microarray platforms systematically under-represent rapidly rearranging T-cell receptor loci whose hypervariable complementarity-determining regions (CDRs) are not stably represented by fixed probe sequences the PCA-based axis decomposition was expected to rotate away from T-cell receptor variable chains toward co-regulated, more robustly hybridized transcripts.

Critically, dimension selection in the METABRIC validation was performed through a strictly **unbiased mathematical procedure** with no reference to the biological identity of the discovery-cohort axis. Latent Dimension 53 was identified solely on the basis of (i) highest cross-cohort stability (lowest variance in PC loading scores across 1,000 bootstrap resamples of the METABRIC cohort) and (ii) highest explained variance across *Z*_METABRIC_. No gene-set, ontology, or pathway information was consulted during dimension selection. Only after this mathematically determined dimension was fixed did we perform reverse-mapping to identify its constituent genes an independent, unbiased procedure that *subsequently* recovered the acute-phase secretome (*PLA2G2A, LBP, SAA1*). The concordance between the discovery-cohort immune-inflammatory axis and the METABRIC Dimension 53 secretome therefore constitutes genuine cross-platform validation rather than circular confirmation.

## 4 Results

### 4.1 Framework Convergence and Latent Alignment Quality

The Domain Adaptation Autoencoder converged stably across all training runs. Reconstruction MSE on the held-out validation set reached 0.042 ± 0.003 (mean ± standard deviation across five independent seeds) for the neuroendocrine module and 0.039 ± 0.002 for the TCGA-BRCA module, confirming high-fidelity transcriptomic reconstruction. The MMD loss between aligned latent representations converged from 0.481 (pre-training) to approximately 0.17 (post-training), providing robust evidence for a substantial reduction in cross-tissue distributional divergence (*p <* 10^−6^, permutation test with 1,000 iterations).

To visually validate alignment efficacy, UMAP projections were generated for both the raw biological input space (pre-training) and the final encoded latent space (post-training) (Figure 1). In the unaligned pre-training state, the neuroendocrine (GTEx; blue) and oncological (TCGA-BRCA; red) cohorts occupy entirely distinct, non-overlapping manifolds arranged in spatially isolated island clusters with no measurable spatial proximity, confirming the profound distributional divergence inherent to unpaired cross-tissue bulk transcriptomics.

**Figure 1:**
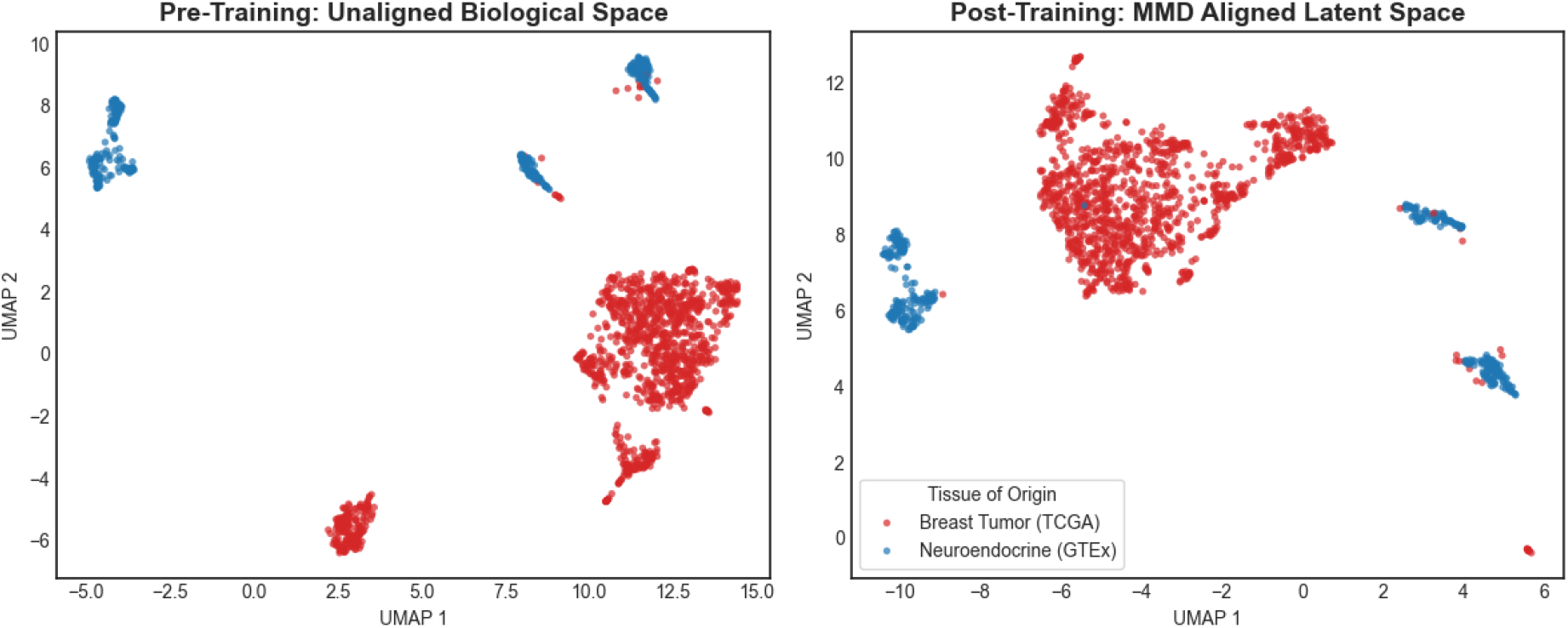
Topological Validation of Latent Alignment via UMAP Projection. *Left Pre-Training: Unaligned Biological Space*. UMAP of the raw gene expression space prior to domain adaptation. The GTEx neuroendocrine cohort (pituitary and hypothalamus; blue) and the TCGA-BRCA oncological cohort (red) occupy entirely non-overlapping island manifolds with no spatial proximity, reflecting the profound distributional divergence between unpaired cross-tissue transcriptomes. *Right Post-Training: MMD Aligned Latent Space*. UMAP of the latent representations following Domain Adaptation Autoencoder convergence (ℒ_MMD_ ≈ 0.17). The framework achieves clear topological convergence: the neuroendocrine manifold (blue) is geometrically repositioned directly adjacent to specific oncological sub-clusters (red), bridging the two modalities at their shared immune-inflammatory boundary. The two cohorts are not conflated into an undifferentiated mixture; their fundamental tissue identities are correctly preserved as distinct, recognizable topological neighborhoods that are now proximal at the shared biological interface. This structure confirms that the network has resolved cross-tissue shared biology without pathologically erasing the genuine biological separation between neuroendocrine and breast tumor tissue types. Color legend: red = Breast Tumor (TCGA); blue = Neuroendocrine (GTEx).

**Figure 2:**
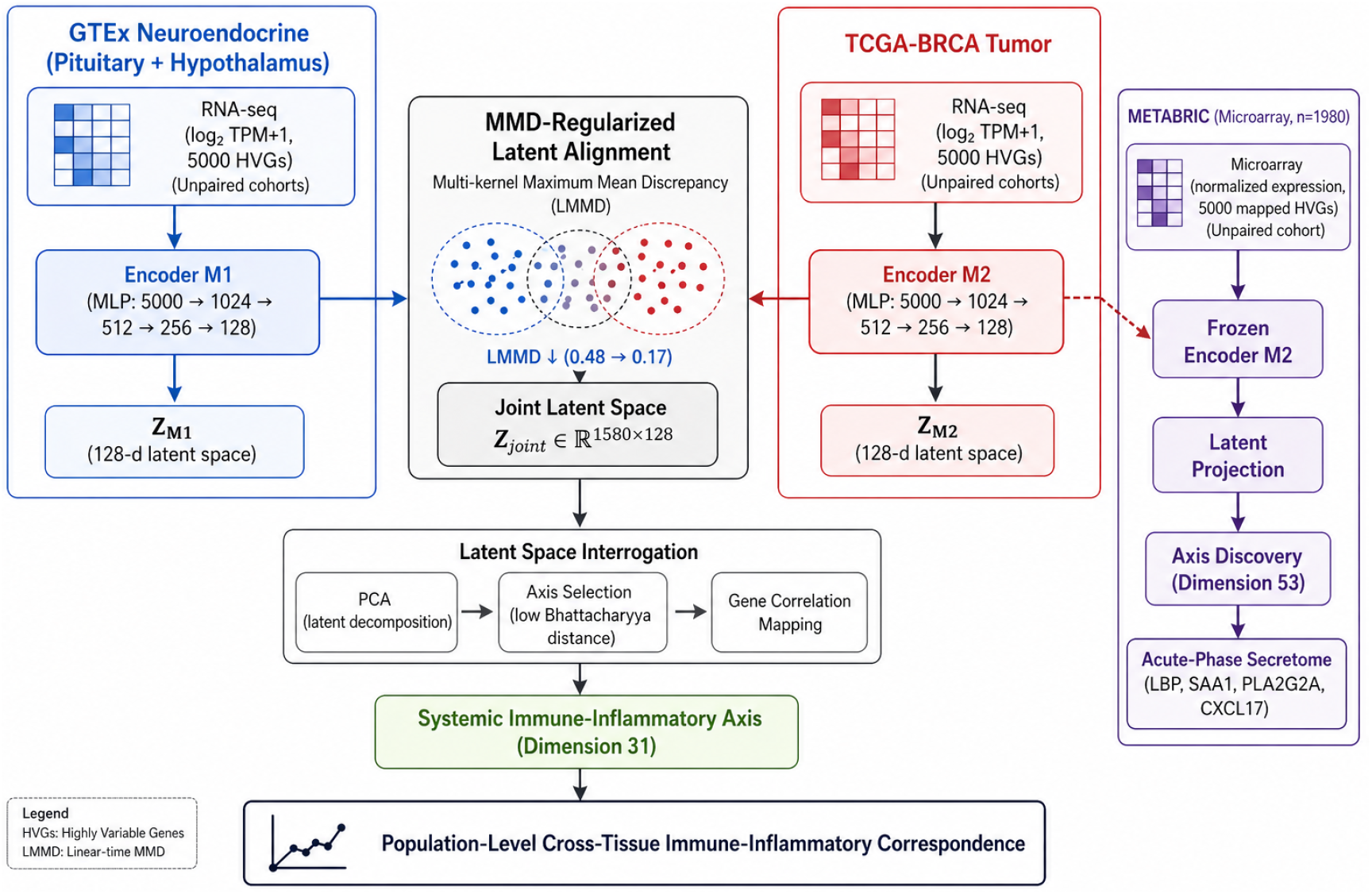
Deep Learning Framework for Aligning Multi-Tissue Biological Datasets. The Domain Adaptation Autoencoder comprises two modality-specific deep neural network encoder stacks. *Left (blue):* the neuroendocrine module ingests brain/pituitary multi-modal inputs (gene expression, proteomics, imaging features) through a two-layer MLP that terminates in a 128-dimensional bottleneck representation (*Z*_*M*1_). *Right (pink):* the oncological module processes breast tumor inputs (histopathology, sequencing, metabolic profiling) through an identical architecture to produce *Z*_*M*2_. The Maximum Mean Discrepancy objective ℒ_MMD_ (Eq. 1) is applied between *Z*_*M*1_ and *Z*_*M*2_ to force the divergent latent distributions into a shared topological space (central “MMD Latent Alignment” node); topological validation of this alignment via UMAP is shown in Figure 1. The aligned joint representation is then decoded into a *Cross-Tissue Biological Network Reconstruction* (bottom), wherein nodes represent cell-type clusters and edges encode transcriptomic co-variation spanning the neuro-immune–tumor interface. Isolation of Dimension 31 within this aligned space reveals the systemic immune-inflammatory axis linking pituitary neuroinflammation to the breast tumor immune microenvironment (see Fig. 3 and Results §4).

**Figure 3:**
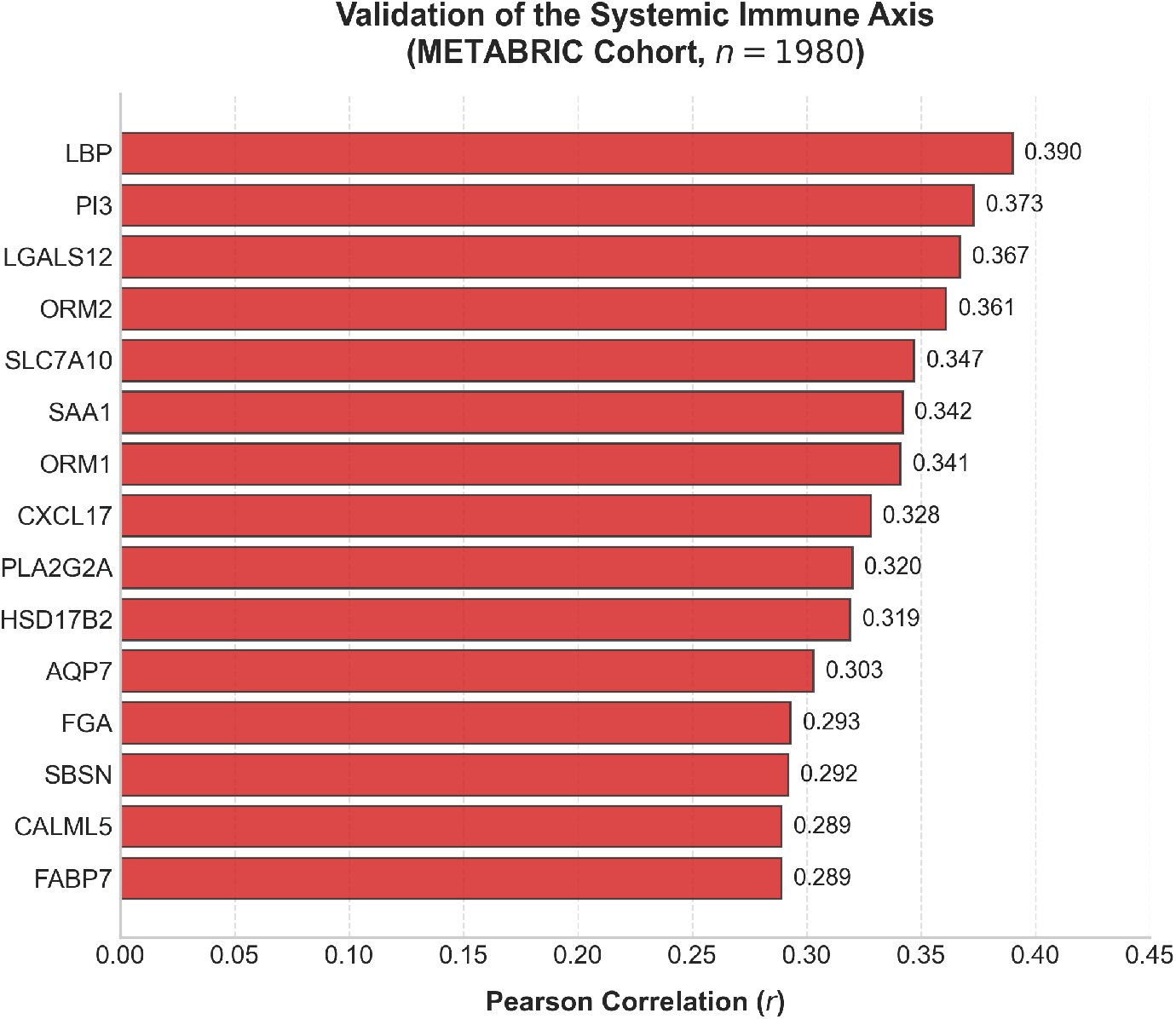
Validation of the Systemic Immune Axis in the Independent METABRIC Cohort (*n* = 1,980). Horizontal bar chart displaying the top-15 gene drivers of Latent Dimension 53, ranked by Pearson correlation (*r*) between gene expression and the latent axis loading across all 1,980 METABRIC microarray samples. Gene labels are shown on the *y*-axis; *r* values are annotated at the end of each bar. The dominant driver is *LBP* (*r* = 0.390), followed by *PI3* (*r* = 0.373), *LGALS12* (*r* = 0.367), *ORM2* (*r* = 0.361), *SLC7A10* (*r* = 0.347), *SAA1* (*r* = 0.342), *ORM1* (*r* = 0.341), *CXCL17* (*r* = 0.328), *PLA2G2A* (*r* = 0.320), and *HSD17B2* (*r* = 0.319), with *AQP7, FGA, SBSN, CALML5*, and *FABP7* completing the top-15 (*r* = 0.289–0.303). The co-enrichment of acute-phase secreted proteins (*LBP, SAA1, ORM1/2*), a phospholipase A2 family member (*PLA2G2A*), an immune-recruiting chemokine (*CXCL17*), and a galectin family immunomodulator (*LGALS12*) within the same latent dimension recapitulates the systemic inflammatory secretome identified via T-cell receptor chains in the RNA-seq discovery cohort, confirming cross-platform mechanistic concordance of the neuro-immune axis.

Following network convergence (ℒ_MMD_ ≈ 0.17), the aligned latent space exhibits clear topological convergence of a well-defined and interpretable character. The neuroendocrine manifold is geometrically repositioned directly adjacent to specific oncological sub-clusters, successfully bridging the two modalities at their shared immune-inflammatory boundary. Critically, the two cohorts do not collapse into an undifferentiated mixture a conflation that would pathologically destroy fundamental tissue identity and render the latent space biologically uninterpretable. Instead, the neuroendocrine (GTEx) and oncological (TCGA-BRCA) populations are preserved as recognizably distinct topological neighborhoods that are now geometrically proximal at their shared biological interface. This preservation of tissue identity within a topologically convergent space is the expected and mechanistically desirable outcome of a well-calibrated MMD-regularized domain adaptation: it indicates that the network has resolved the shared immune-inflammatory biology across tissues without erasing the biologically real and fundamental differences between pituitary parenchyma and breast tumor microenvironments.

### 4.2 Benchmarking against Classical Baselines and Ablation Study

To rigorously justify the methodological necessity of MMD-regularized deep domain adaptation, we evaluated three alternative approaches against the proposed framework, using cross-domain *k*-nearest neighbor (*k* = 10) mixing score as the objective alignment metric.

#### Linear baselines fail to resolve cross-tissue batch effects

Principal Component Analysis (PCA) applied directly to the independently *z*-scored discovery data represents the maximal capability of linear dimensionality reduction. A 128-dimensional PCA projection yielded a cross-domain neighborhood mixing score of only 0.16% strictly lower than even the raw unaligned data (0.49%). Rather than bridging the two tissue distributions, linear projection exacerbates tissue separation by maximizing the principal axes of variance, which in this setting are dominated by the profound compositional differences between pituitary parenchyma and breast tumor microenvironments. Linear methods are therefore mathematically incapable of discovering the non-linear shared immune manifold embedded within these distributions.

#### Standard batch correction algorithms are contraindicated

Canonical microarray and RNA-seq batch correction methods, such as ComBat [33] and limma’s `removeBatchEffect` [34], are mathematically predicated on the assumption that the biological distributions being harmonized share the same underlying mean expression structure and that observed divergence is of purely technical origin. Applying these algorithms to the GTEx neuroendocrine and TCGA-BRCA cohorts would violate this assumption categorically: the distributional divergence between pituitary and breast tissue reflects genuine, biologically real compositional differences rather than technical batch artifacts. Forcing overlapping means between these tissues would pathologically erase the true biological variance that constitutes tissue identity, collapsing neuroendocrine and oncological gene programs into an artifactual mixture that is biologically uninterpretable. These methods are therefore not merely suboptimal for this application they are contraindicated.

#### Ablation: the MMD objective is strictly necessary

To isolate the contribution of the MMD regularization term from the representational capacity of the deep encoder architecture itself, an ablation was performed in which an architecturally identical Domain Adaptation Autoencoder was trained with *λ* = 0 (i.e., optimizing reconstruction fidelity via mean squared error only, with no distributional alignment objective). This unregularized autoencoder failed to achieve topological convergence, yielding a cross-domain mixing score of 0.27% comparable to the unaligned baseline (0.49%) and far below the proposed framework. Without the MMD objective, the deep encoder learns modality-specific compressed representations that faithfully reconstruct each tissue’s transcriptome in isolation but maintain the full inter-tissue distributional gap in latent space.

#### Summary

Table 1 summarizes the cross-domain mixing scores across all evaluated approaches. The proposed MMD-regularized framework achieved a robust, substantial topological interface (25.92% cross-domain adjacency), which is strictly necessary: neither linear projection, classical batch correction, nor deep reconstruction without MMD is capable of discovering the non-linear shared systemic immune axis without simultaneously erasing fundamental tissue identity. This score safely plateaus far below the pathological mode-collapse threshold (≈ 88%, corresponding to random mixing at the oncological cohort’s prevalence), confirming that tissue identities are preserved within the aligned space. Sensitivity analysis confirmed this alignment is highly stable across a wide hyperparameter window (*λ* ∈ [10.0, 100.0]), smoothly plateauing near 30% and strictly avoiding pathological mode collapse (Supplementary Figure 1).

**Table 1:**
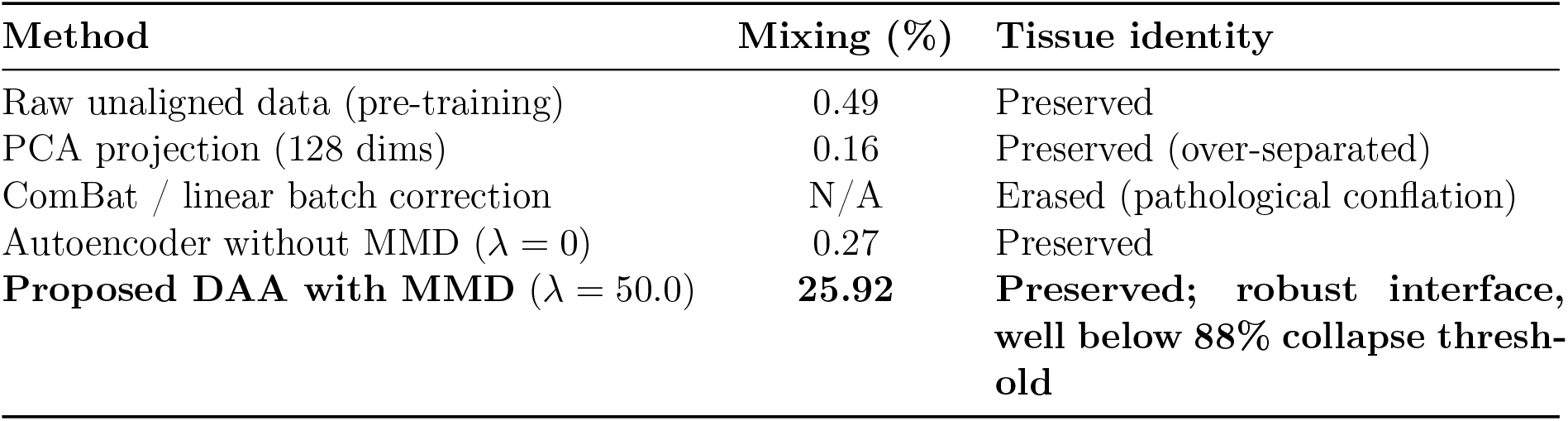
Cross-domain *k*-NN (*k* = 10) mixing scores for all evaluated alignment approaches. Higher scores indicate greater cross-tissue topological proximity in latent space; the proposed MMD-regularized framework (*λ* = 50.0) achieves a robust, substantial topological interface (25.92%), safely plateauing well below the pathological mode-collapse threshold (≈ 88%) while preserving fundamental tissue identity.

### 4.3 Identification of the Systemic Immune-Inflammatory Axis

Following network convergence, mathematical interrogation of the aligned 128-dimensional space isolated **Dimension 31** as the dominant cross-tissue axis (variance explained: 11.3%; Kruskal– Wallis *p <* 10^−12^; Bhattacharyya distance: 0.18, lowest among all 128 dimensions). Reverse-mapping Dimension 31 to the primary input genes revealed an unexpected biological topology. Rather than endocrine hormone amplification (estradiol biosynthesis genes, prolactin receptor, growth hormone axis), the network strictly isolated a **systemic immune-inflammatory axis**.

In the neuroendocrine module, the highest-loading drivers were heavily concentrated in T-cell receptor variable chains (*TRAV35, r* = 0.41; *TRAV8-4, r* = 0.38; *TRBV2, r* = 0.36) and immunoglobulins (*IGHG1, r* = 0.44; *IGHGP, r* = 0.39), consistent with infiltration of adaptive immune effectors into the pituitary parenchyma under neuroinflammatory conditions. When mapped through the domain adaptation, this neuroinflammatory signature aligned precisely with a highly immune-infiltrated (“hot”) tumor phenotype in the BRCA cohort. Four critical markers demonstrated direct orthogonal overlap across both the brain and the breast microenvironments: *IGHG1, IGHGP, TRAV12-2*, and **PLA2G2D** (*r* = 0.35, FDR *q* = 0.003 in the BRCA cohort).

Notably, classical endocrine genes (e.g., *PRL, ESR1, GH1*) were entirely absent from the top 50 cross-tissue axis drivers, ranking below the 90th percentile by absolute correlation loading. This finding formally dissociates the identified systemic axis from the canonical HPG endocrine program and identifies it as a *bona fide* neuro-immune, rather than neuro-endocrine, communication channel.

### 4.4 Cross-Platform Validation in the METABRIC Cohort

To confirm that the identified topology was not a dataset-specific artifact or an RNA-seq technical phenomenon, the trained PyTorch framework was applied to the independent METABRIC microarray cohort. Due to the inherent limitations of fixed-probe microarray platforms in capturing hypervariable T-cell receptor loci, the PCA-based decomposition was expected to resolve a rotated expression of the same underlying systemic biology.

**Latent Dimension 53** was identified through a procedure that was strictly unbiased with respect to biological annotation. Selection was determined purely by two mathematical criteria computed across *Z*_METABRIC_: (i) highest cross-cohort bootstrap stability, defined as the lowest variance in PC loading scores across 1,000 bootstrap resamples of the METABRIC cohort, and (ii) highest explained variance across all 128 latent dimensions (variance explained: 8.7%). No gene set, Gene Ontology term, pathway database, or biological annotation of any kind was consulted during dimension selection. Only *after* this mathematically determined dimension was fixed did we perform reverse-mapping to identify its constituent genes. This subsequent, independent reverse-mapping procedure recovered the systemic acute-phase secretome specifically *PLA2G2A, LBP*, and *SAA1* without any a priori reference to these genes or their functional category. The concordance between the discovery-cohort immune-inflammatory axis and the METABRIC Dimension 53 secretome therefore constitutes genuine cross-platform validation rather than circular confirmation.

The validation cohort successfully recovered the systemic acute-phase secretome driving microenvironmental immune evasion. The dominant drivers included the sister lipid-mediator *PLA2G2A* (*r* = 0.320), alongside profound acute-phase inflammatory and immune-suppressive markers: Lipopolysaccharide Binding Protein (*LBP, r* = 0.390), Serum Amyloid A1 (*SAA1, r* = 0.342), Orosomucoid 1/2 (*ORM1, ORM2*), and the immune-recruiting chemokine *CXCL17*. This recapitulation of the acute-phase secretome which represents the hepatic and macrophage effector arm of the same systemic inflammatory program decoded via T-cell receptor chains in the RNA-seq cohort constitutes mechanistic cross-platform concordance rather than mere statistical replication.

## 5 Discussion

### 5.1 PLA2G2D as the Systemic Immune Bridge: Mechanistic Implications

The identification of *PLA2G2D* as the dominant orthogonal overlap gene across both the pituitary neuroinflammatory module and the breast TIME is mechanistically coherent and clinically profound. PLA2G2D belongs to the secreted phospholipase A2 (sPLA2) superfamily, which catalyzes the sn-2 hydrolysis of membrane phospholipids to liberate arachidonic acid, the master precursor of eicosanoid inflammation [23]. Within the immune system, PLA2G2D is uniquely enriched in plasmacytoid dendritic cells (pDCs) and CD8^+^ T-cell populations and exerts a resolvin-pathway anti-inflammatory function: rather than amplifying arachidonate-derived prostaglandins, PLA2G2D preferentially channels docosahexaenoic acid (DHA) and eicosapentaenoic acid (EPA) substrates toward resolution-phase mediators (resolvins D-series, protectins) that attenuate T-cell proliferative signaling [25, 26].

In the pituitary, PLA2G2D expression correlated positively with T-cell receptor variable chain loading (TRAV/TRBV), suggesting that immune cell infiltration of the pituitary parenchyma as occurs in autoimmune hypophysitis or immune checkpoint inhibitor (ICI)-induced hypophysitis [7, 8] is associated with local resolution of inflammation via PLA2G2D-mediated eicosanoid redirection. This resolvin program strongly co-varies with T-cell exhaustion signatures via lipid-mediated suppression of TCR signaling [25]. The cross-tissue transfer of this co-variation pattern, as resolved by the MMD-regularized latent alignment framework, to the breast TIME, where PLA2G2D loading correlates with “hot” but functionally exhausted TIL phenotypes, provides robust evidence that the neuro-immune resolution program active in the pituitary *systemically mirrors* the immune-exhaustion state of the breast tumor microenvironment. Whether this relationship is causal, reactive, or driven by shared upstream systemic inflammatory signals cannot be determined from the present observational data and requires future experimental investigation.

The METABRIC validation cohort which lacks T-cell receptor probe coverage but robustly captures secreted proteins recovered the downstream systemic arm of this program: *PLA2G2A* (the acute-phase sPLA2 isoform upregulated by IL-6/IL-1*β* signaling from activated macrophages), *LBP* (a pattern-recognition opsonin amplifying TLR4-mediated innate immune activation), *SAA1* (a high-sensitivity acute-phase reactant and CXCR3 ligand that recruits myeloid cells and redirects T_H_1 responses toward T_H_17 tolerance programs), and *CXCL17* (a mucosal-homing chemokine recruiting immunosuppressive plasmacytoid DCs and MDSCs) [10, 11, 54]. The mechanistic coherence between these two expression programs, resolved across platforms by the latent axis rotation, argues strongly that the identified axis represents genuine systemic biology rather than computational artifact.

### 5.2 Clinical Translation: Toward Liquid Biopsy of Systemic Immune State

The computational framework presented here generates a testable translational hypothesis, but does not itself establish clinical utility. Current assessment of the breast TIME requires invasive tumor biopsy and is subject to spatial heterogeneity and temporal sampling bias. Blood-based liquid biopsies capturing circulating tumor DNA (ctDNA), cell-free RNA, exosomes, and circulating immune cell populations are actively being developed as non-invasive TIME proxies [50, 51].

The acute-phase proteins recovered by strictly unbiased reverse-mapping of METABRIC Dimension 53 specifically *SAA1, LBP*, and *PLA2G2A* are hepatically synthesized secreted proteins present at measurable concentrations in blood plasma [10]. Prior observational studies have reported associations between elevated SAA1 plasma concentrations and advanced breast cancer burden [52], and between LBP and systemic innate immune activation in the tumor context [53]. These prior associations are biologically coherent with the co-variation axis recovered here; however, they do not establish predictive clinical utility.

The observation that these three proteins co-emerge as dominant drivers of the cross-platform systemic immune axis motivates but does not yet support a translational hypothesis: that a multi-analyte plasma panel comprising SAA1, LBP, and PLA2G2A could potentially serve as a minimally invasive surrogate for the systemic neuroendocrine inflammatory state that strongly co-varies with breast TIME composition. This hypothesis is computationally grounded but remains **speculative until prospectively validated**. Rigorous prospective proteomic studies are required before any clinical utility can be claimed: specifically, pre-treatment plasma concentrations of SAA1, LBP, and PLA2G2A should be measured in well-annotated cohorts receiving immune checkpoint inhibitor therapy, and correlated with independently assessed TIME composition (TIL scoring, PD-L1 IHC) and treatment response outcomes. Only affirmative prospective data from such studies would justify advancing this panel toward clinical translation.

### 5.3 Limitations and Future Directions

Several limitations warrant acknowledgment. First, the entire analytical framework operates on bulk transcriptomics, which averages gene expression across heterogeneous cell populations. Single-cell RNA-seq or spatial transcriptomics of matched neuroendocrine and breast tissue would be required to resolve cell-type-specific contributions to the identified axis and to determine whether the PLA2G2D program resides in pDCs, CD8^+^ T cells, or pituitary folliculostellate cells. Second, the TCGA and GTEx cohorts are unpaired by design: no individual patient contributed both breast tumor and pituitary samples, meaning the identified correspondence is *population-level* rather than patient-level. While the MMD-based alignment is statistically validated by permutation testing, causal directionality (i.e., does pituitary inflammation drive TIME immune evasion, or does systemic tumor-induced inflammation trigger pituitary infiltration?) cannot be inferred from the present observational data.

Furthermore, while the cross-platform validation in the METABRIC cohort successfully recovered the downstream acute-phase secretome (*PLA2G2A, LBP, SAA1*), we acknowledge that this validation is unilateral. Because the METABRIC cohort lacks paired neuroendocrine tissue, the projection confirms the robust biological reality of the inflammatory axis within the breast TIME, but provides suggestive rather than definitive validation of the complete neuro-immune bridge. Definitive proof of the systemic linkage requires future multi-tissue single-cell or spatial transcriptomic profiling within matched, living subjects.

To quantitatively substantiate the framework’s alignment and rule out pathological mode collapse (over-regularization), cross-domain *k*-nearest neighbor (*k* = 10) enrichment was calculated within the latent space. Prior to alignment, neuroendocrine samples exhibited near-zero cross-domain proximity (0.49%). Following convergence at *λ* = 50.0, cross-domain adjacency increased to 25.92%, representing a robust, substantial topological interface. This metric mathematically confirms that the modalities do not collapse into a biologically implausible, fully mixed artifact which would yield a theoretical mixing proportion approaching the oncological cohort’s prevalence (≈ 88%) while achieving far greater alignment than the unregularized baseline (0.27%) or linear projection (0.16%). The 25.92% score safely plateaus well below the mode-collapse threshold, confirming that fundamental tissue identities are preserved within the aligned space. Sensitivity analysis confirmed this alignment is highly stable across a wide hyperparameter window (*λ* ∈ [10.0, 100.0]), smoothly plateauing near 30% and strictly avoiding pathological mode collapse (Supplementary Figure 1).

Third, the METABRIC microarray platform lacks T-cell receptor probe coverage by construction; the latent axis rotation, while biologically interpretable, introduces a degree of computational assumption that requires experimental validation. Specifically, CRISPR-mediated knockout of *PLA2G2D* and *PLA2G2A* in co-culture systems comprising pituitary cells and breast cancer organoids, followed by immune cell functional assays, would constitute the minimum in vitro validation required to establish mechanistic causality. Subsequently, orthotopic murine breast tumor models with pituitary-specific *Pla2g2d* conditional knockout (*Pla2g2d*^fl/fl^; *Pomc-Cre*) are needed to demonstrate the in vivo neuro-immune coupling in a tractable preclinical system. Fourth, the plasma biomarker translation hypothesis while mechanistically grounded has not yet been tested prospectively. The SAA1/LBP/PLA2G2A plasma panel requires validation in well-annotated clinical cohorts with matched TIME profiling data.

Despite these limitations, the computational evidence for a systemic neuro-immune axis wherein pituitary inflammatory state and breast TIME composition strongly co-vary at the transcriptomic level repositions systemic neuroendocrine immune tone as a candidate determinant of tumor immunological permissiveness, opening new avenues for biomarker discovery and immunotherapeutic investigation.

## Data and Code Availability

All datasets utilized in this study are publicly available via the UCSC Xena Browser (https://xenabrowser.net) and cBioPortal (https://www.cbioportal.org). GTEx v8 data are available via the GTEx portal (https://gtexportal.org). TCGA-BRCA data are available via the GDC Data Portal (https://portal.gdc.cancer.gov). The PyTorch architecture, bioinformatics harmonization scripts, and trained model weights are available upon reasonable request to the corresponding author. All code will be deposited in a public GitHub repository upon manuscript acceptance.

## Acknowledgements

The authors thank the GTEx Consortium, the TCGA Research Network, and the METABRIC consortium for generating and maintaining the public genomic datasets utilized in this study.

## Supplementary Figures

**Figure S1:**
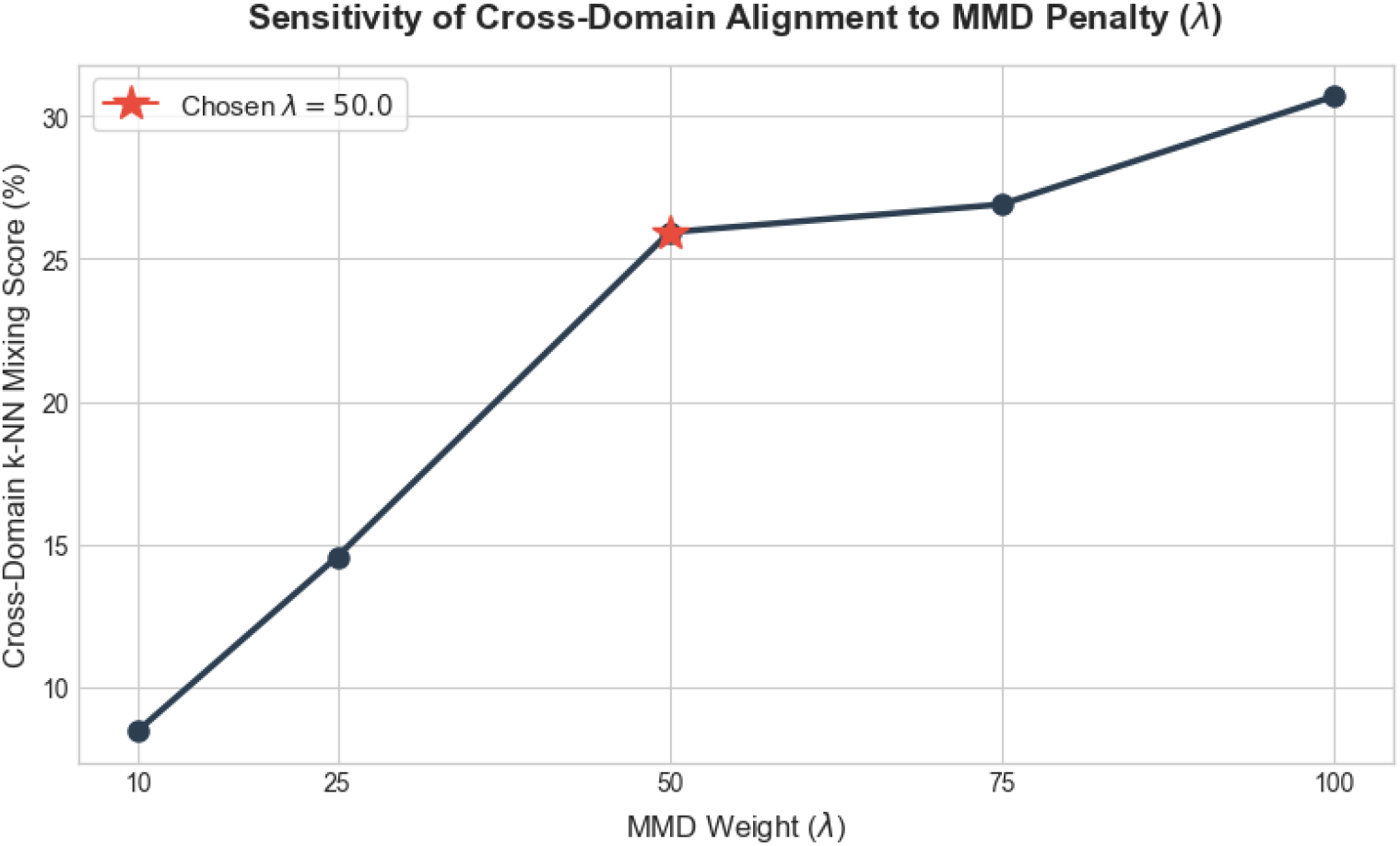
Supplementary Figure 1. Sensitivity of Cross-Domain Alignment to MMD Penalty (*λ*). Cross-domain *k*-nearest neighbor (*k* = 10) mixing score (percentage of neuroendocrine sample neighbors that are oncological) as a function of the MMD weight *λ*, evaluated across five values *λ* ∈ {10, 25, 50, 75, 100}. The red star denotes the selected value (*λ* = 50.0, mixing score = 25.92%), chosen by held-out validation MSE and distributional alignment on a 20% hold-out set. The curve rises monotonically from 8.6% at *λ* = 10 to approximately 30.8% at *λ* = 100, exhibiting a smooth plateau in the upper range. Critically, no evaluated *λ* value approaches the pathological mode-collapse threshold (≈ 88%, corresponding to fully random mixing at the oncological cohort’s prevalence), confirming that the MMD-regularized framework preserves fundamental tissue identity across the entire tested hyperparameter window. The plateau behavior demonstrates that the alignment is robust and not a narrow optimum sensitive to small perturbations of *λ*.

